# Multi-Modal Large Language Model Enables All-Purpose Prediction of Drug Mechanisms and Properties

**DOI:** 10.1101/2024.09.29.615524

**Authors:** Youwei Liang, Ruiyi Zhang, Yongce Li, Mingjia Huo, Zinnia Ma, Digvijay Singh, Chengzhan Gao, Hamidreza Rahmani, Satvik Bandi, Li Zhang, Robert Weinreb, Atul Malhotra, Danielle A. Grotjahn, Linda Awdishu, Trey Ideker, Michael Gilson, Pengtao Xie

**Affiliations:** Department of Electrical and Computer Engineering, University of California San Diego, La Jolla, California 92093, USA; Department of Bioengineering, University of California San Diego, La Jolla, California 92093, USA; School of Biological Sciences, University of California San Diego, La Jolla, CA 92093, USA; Department of Integrative Structural and Computational Biology, The Scripps Research Institute, La Jolla, CA 92037, USA; Viterbi Family Department of Ophthalmology, University of California San Diego, La Jolla, CA 92093, USA; Department of Medicine, University of California San Diego, La Jolla, CA 92093, USA; School of Pharmacy and Pharmaceutical Science, University of California San Diego, La Jolla, CA 92093, USA

**Keywords:** Drug mechanism prediction, drug property prediction, multimodal large language model, graph neural network

## Abstract

Accurately predicting the mechanisms and properties of potential drug molecules is essential for advancing drug discovery. However, traditional methods often require the development of specialized models for each specific prediction task, resulting in inefficiencies in both model training and integration into workflows. Moreover, these approaches are typically limited to predicting pharmaceutical attributes represented as discrete categories, and struggle with predicting complex attributes that are best described in free-form texts. To address these challenges, we introduce DrugChat, a multi-modal large language model (LLM) designed to provide comprehensive predictions of molecule mechanisms and properties within a unified framework. DrugChat analyzes the structure of an input molecule along with users’ queries to generate comprehensive, free-form predictions on drug indications, pharmacodynamics, and mechanisms of action. Moreover, DrugChat supports multi-turn dialogues with users, facilitating interactive and in-depth exploration of the same molecule. Our extensive evaluation, including assessments by human experts, demonstrates that DrugChat significantly outperforms GPT-4 and other leading LLMs in generating accurate free-form predictions, and exceeds state-of-the-art specialized prediction models.

## Introduction

Accurate prediction of the mechanisms and properties of potential drug molecules is crucial for advancing pharmaceutical research and facilitating drug discovery (1). As the development of new drugs becomes increasingly complex, the demand for reliable computational models that can predict these attributes has grown exponentially (2). Deep learning models have emerged as powerful tools for addressing this challenge, thanks to their ability in analyzing vast amounts of data and uncovering complex patterns (3–6).

However, despite their potential, existing approaches in this domain typically involve developing specialized models tailored to specific prediction tasks, such as predicting pharmacokinetics (7), toxicity (8), or molecular binding affinities (9, 10). While these models have achieved notable successes, their task-specific nature imposes certain limitations. First of all, each model requires extensive training on large datasets specific to the prediction task, which leads to inefficiencies in both computational resources and time. Additionally, integrating predictions from multiple models often requires complex pipelines that further complicate the drug discovery process (11, 12). Moreover, existing approaches are limited to predicting relatively simple attributes represented as discrete categories (13, 14), and they lack the capability to accurately predict more complex, nuanced aspects of drug molecules, such as indications, pharmacodynamics, and mechanisms of action, which are best described in free-form text.

To address these challenges, we introduce DrugChat, a multi-modal large language model (LLM) designed for the comprehensive prediction of drug mechanisms and properties. DrugChat integrates multiple modalities, including molecular structures, molecular images, and texts, to provide a holistic understanding of potential drug molecules. Users can upload a molecule and interact with DrugChat by asking various questions in natural language, referred to as *prompts*, to gain insights about the molecule. DrugChat leverages a graph neural network (15) and a convolutional neural network (16) to effectively capture and interpret the structure of this molecule. These interpretations are seamlessly integrated into an LLM (17), which generates detailed, contextually relevant responses in free-form text to users’ questions.

Unlike existing approaches (18–22), DrugChat handles a wide range of prediction tasks within a single framework, thereby eliminating the need for multiple specialized models. This not only reduces the computational and time costs associated with training and maintaining separate models but also simplifies the integration of various predictions into a cohesive workflow (23). Furthermore, DrugChat’s ability to generate predictions in free-form text allows for a richer and more nuanced understanding of drug molecules, particularly for complex attributes like pharmacodynamics, indications, and mechanisms of action that are challenging to express in discrete categories. By integrating and contextualizing these interrelated molecule attributes, DrugChat offers a holistic view that enhances predictive accuracy. DrugChat’s multi-modal capabilities further boost its predictive power by incorporating both structural and visual modalities of molecules, ensuring that no critical aspect of a molecule’s characteristics is overlooked. This comprehensive approach enables DrugChat to provide more insightful and informed decision-making in drug development. Additionally, DrugChat’s interactive, multi-turn dialogue system allows users to explore drug molecules in depth, refining their prompts based on DrugChat’s initial responses and thereby gaining a more detailed understanding of the molecules. This adaptability and depth of interaction represent important improvements over the static, single-output nature of traditional models (24, 25).

## Results

### DrugChat overview

Examples of DrugChat’s usage are shown in Fig. 1a. It accepts a compound molecule along with a user’s prompt as input, and generates a textual prediction. For example, when provided with a molecule and the prompt ‘what is its mechanism of action?’, DrugChat generates a prediction such as: ‘it stimulates neurons to release or maintain high levels of a particular group of neurotransmitters…’.

**Fig. 1.**
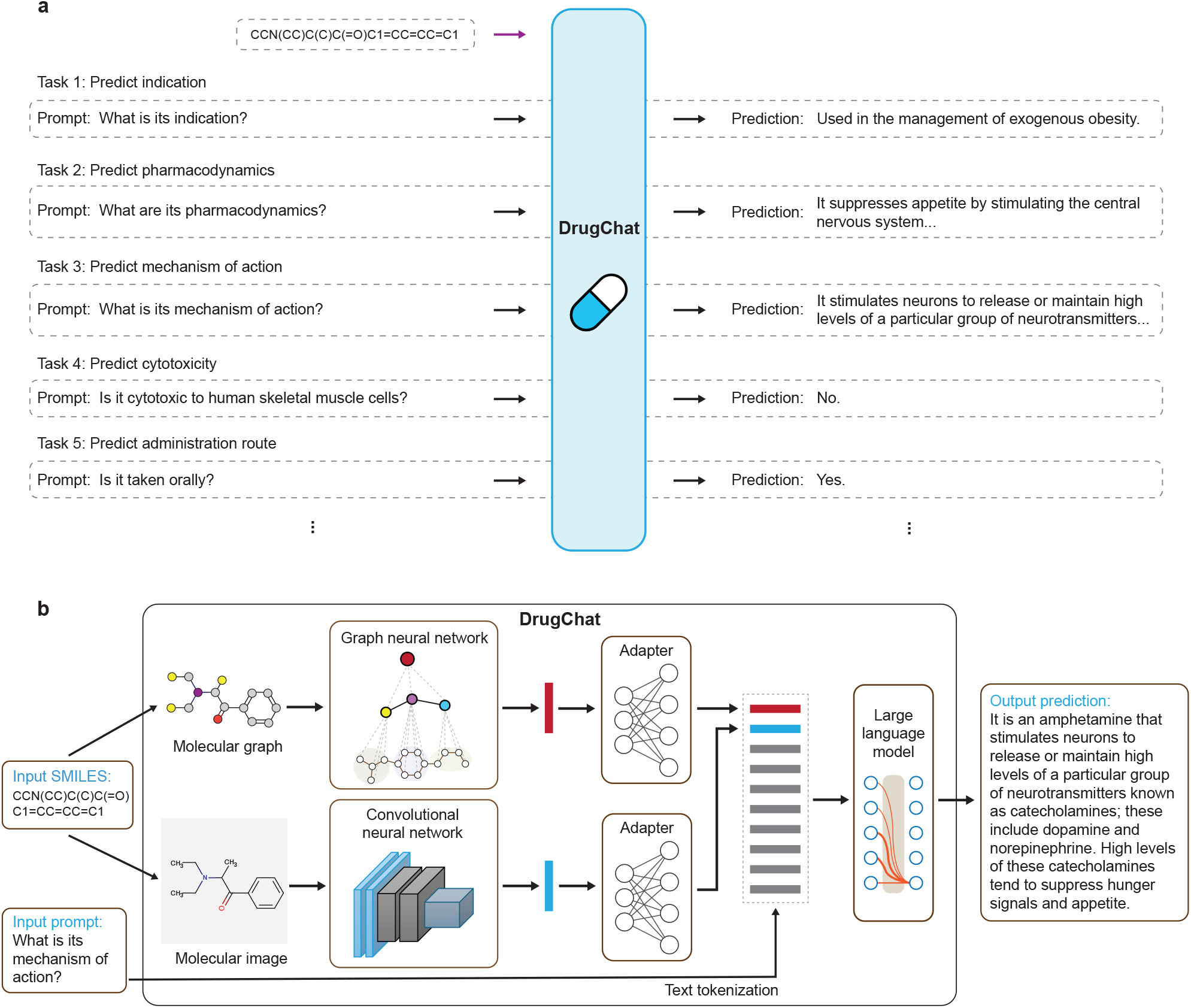
DrugChat is a multi-modal LLM capable of predicting drug attributes in either free-form text or as discrete categories. **a**, DrugChat facilitates versatile predictions of drug attributes, enabling users to submit queries through flexible natural language (known as prompts). By using task-specific prompts, DrugChat can perform a wide range of prediction tasks within a unified framework, without requiring changes to model parameters. **b**, Model architecture of DrugChat. It takes the SMILES representation of a molecule along with a prompt as inputs and generates a prediction in natural language. It comprises two molecular encoders - a graph neural network and a convolutional neural network - that learn representation vectors for the molecular graph and image derived from the SMILES input, two adapters that transform these representations into a format compatible with LLMs, and an LLM that generates a prediction based on the molecular representations and the prompt.

The core components of DrugChat include molecule encoder networks, a large language model (LLM) (26, 27), and two adapters that seamlessly integrate these encoders with the LLM (Fig. 1b). The input molecule is initially represented using a SMILES (Simplified Molecular Input Line Entry System) string. DrugChat converts the SMILES string into two forms: a structural representation as a molecular graph and a visual representation as a molecular image. For the molecular graph, a graph neural network (GNN) (25) based encoder, pretrained on two million unlabeled molecules from the ZINC15 database (28) using self-supervised learning (25), processes the graph by first converting nodes and edges into distinct representation vectors based on their types. The GNN then iteratively updates these node and edge representations by aggregating information from each node’s neighbors (29, 30). Finally, a pooling operation (31) is applied to summarize the node representations into a single representation vector of the entire molecular graph. The molecular image is encoded by a convolutional neural network (CNN) (16), specifically the ImageMol model (24), which was pretrained on images of ten million unlabelled bioactive molecules from the PubChem dataset (32) using self-supervised learning (24). The CNN applies multiple layers of 2D convolutional filters, which function as matching templates, to ‘scan’ the image, producing a feature map (33). This feature map is then condensed into a representation vector of the entire molecular image through a pooling operation (34). The adapters, implemented as multi-layer perceptrons (MLPs), are used to transform the representation vectors of the molecule’s graph and image into a unified vector, termed the *molecule token*, which is compatible with the LLM’s latent representation space, thereby enabling the LLM to interpret it. Concurrently, the input prompt is decomposed into a sequence of language tokens, each represented as a vector within the LLM’s latent space. The molecule token is integrated into the language token sequence, which is then fed into the LLM, specifically Vicuna-13B (35). The LLM then generates new language tokens sequentially through autoregressive decoding (36), ultimately producing the prediction.

To train DrugChat, we curated a comprehensive dataset from three databases, including ChEMBL (37), Pub-Chem (32), and DrugBank (38). After filtering and removing duplicates, we collected approximately 14,000 unique drug molecules. These molecules encompass both approved and experimental drugs, covering a wide range of compound categories (Extended Data Fig. 2). Each drug molecule’s attributes were meticulously annotated by experts, including complex attributes described in free-form text, such as indications, pharmacodynamics, and mechanisms, as well as simpler attributes represented by discrete categories, such as cytotoxicity and administration routes. From these annotated drug molecules, we curated 91,365 (molecule, prompt, answer) triplets for training DrugChat. In these triplets, the molecule and prompt serve as inputs to DrugChat, while the answer represents the target output of DrugChat. The answers vary from free-form texts that describe indications, pharmacodynamics, and mechanisms, to structured responses like a binary ‘yes/no’ to questions such as ‘can it be taken orally?’. We divided the drug entries from ChEMBL and DrugBank into training (80%), validation (10%), and test (10%) sets, while the entire PubChem dataset was used exclusively for training. We trained model weights by minimizing the negative log-likelihood (39) between DrugChat’s predictions and the ground truth answers in the training sets.

### DrugChat generates precise, free-form predictions for drug indications, pharmacodynamics, and mechanisms of action

Drug indications refer to the diseases that a molecule is intended to treat or its potential applications in clinical trials. Pharmacodynamics describes the biochemical and physiological effects that a molecule exerts on the body, while the mechanism of action explains how the molecule produces these effects. These attributes are typically described in free-form text within the literature. We employed DrugChat to generate free-form textual predictions for these attributes using specific prompts: ‘what is its indication?’, ‘what are its pharmacodynamics?’, and ‘what is its mechanism of action?’. To evaluate DrugChat’s performance, we randomly selected 572 test drug molecules from DrugBank, ensuring that none were included in the training set. We compared DrugChat’s predictions to those of GPT-4 (27), a flagship LLM, which also uses molecules’ SMILES as input. Additionally, each drug molecule in DrugBank is accompanied by an expert-written overview that summarizes essential information about the molecule. We assessed the ability of DrugChat and GPT-4 to generate overviews for the test molecules.

We conducted a human evaluation in which pharmaceutical science experts assessed the accuracy of the model’s predictions by comparing them against the ground truth annotations provided by DrugBank, without knowing which model generated the predictions. The evaluation was based on a scoring rubric, assigning scores of 2, 1, or 0 for each prediction, corresponding to ‘correct’, ‘partially Correct’, and ‘incorrect’, respectively (Methods). Fig. 3 provides examples illustrating how these scores were assigned on some molecules. DrugChat achieved average human assessment scores of 1.05 for indication prediction, 0.94 for pharmacodynamics, 0.8 for mechanism of action, and 0.92 for overview, significantly outperforming GPT-4, which scored 0.38, 0.82, 0.45, and 0.46 respectively (Fig. 2a). The superiority of DrugChat over GPT-4 is also reflected in the distribution of scores (Fig. 2b). For instance, in the task of indication prediction, 42.9% of DrugChat’s predictions were rated as correct, 19% as partially correct, and 38.1% as incorrect. In contrast, GPT-4’s predictions received ratings of 14.3% correct, 9.5% partially correct, and 76.2% incorrect. As another example, for action mechanism prediction, DrugChat’s predictions were rated as 25% correct, 30% partially correct, and 45% incorrect, while GPT-4 received ratings of 15% correct, 15% partially correct, and 70% incorrect. In addition to assigning absolute scores, the human evaluators also performed relative comparisons by assessing which prediction - whether from DrugChat or GPT-4 - was superior for each molecule (Fig. 2c). DrugChat outperformed GPT-4 in most cases. Specifically, DrugChat generated better predictions for 52.4% of molecules in terms of indication, 41.2% in pharmacodynamics, 50% in mechanism of action, and 47.2% in overview. In contrast, GPT-4 outperformed DrugChat in only 14.3%, 35.3%, 30%, and 25% of the cases, respectively. The remaining comparisons resulted in ties.

**Fig. 2.**
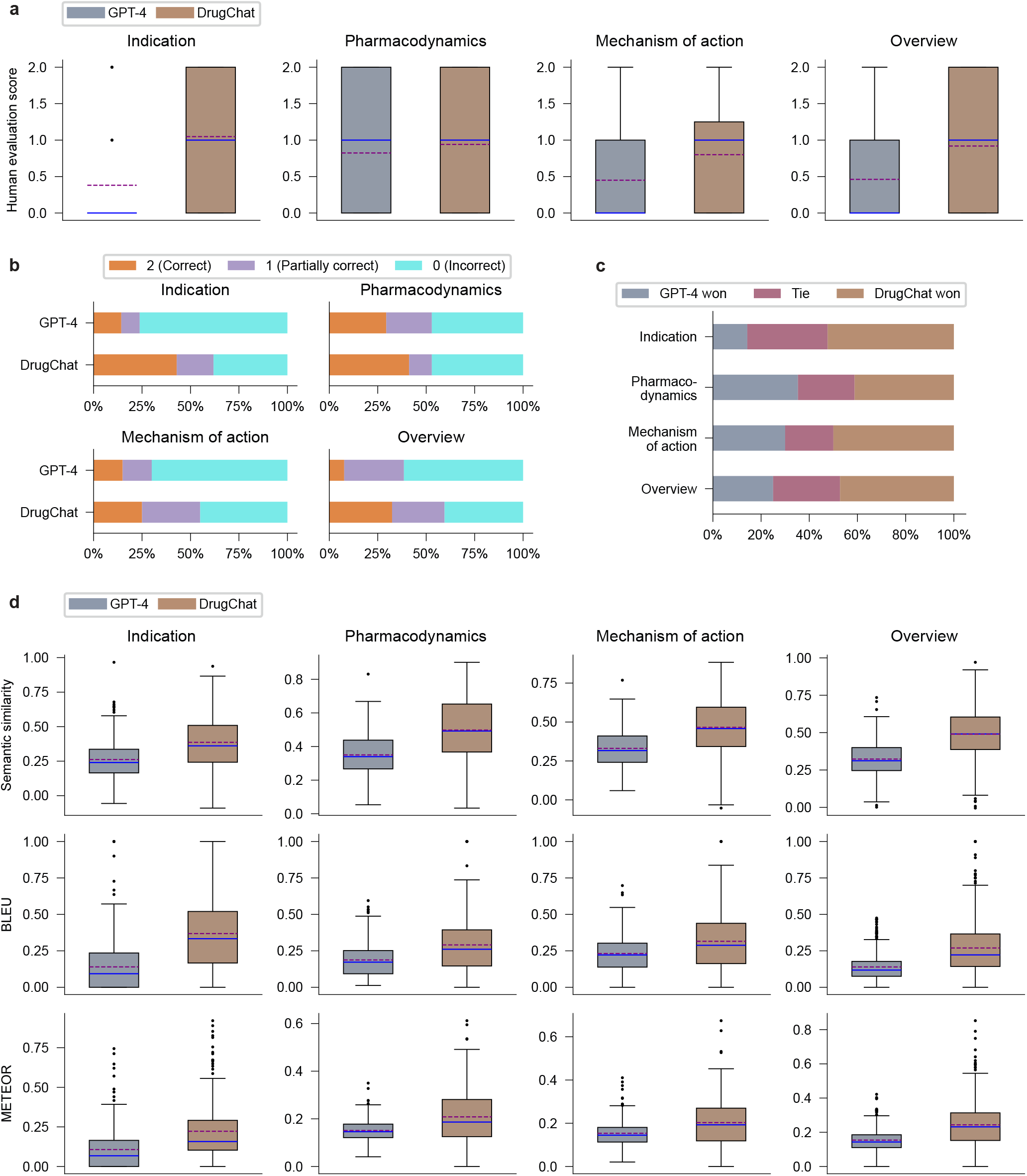
DrugChat accurately predicts drug indications, pharmacodynamics, and mechanisms of action expressed in textual descriptions and significantly outperforms GPT-4, a flagship LLM. **a-c**, Human expert evaluations demonstrated DrugChat’s significantly superior performance over GPT-4, as shown by the average expert scores (**a**), the distribution of scores (**b**), and human evaluator preferences between DrugChat and GPT-4 predictions (**c**). Experts rated the predictions on a scale from 0 to 2, with 2 indicating fully correct predictions, 1 for partially correct, and 0 for incorrect. Both DrugChat and GPT-4 used SMILES representations of molecules as input. **d**, In automated evaluations - measured by semantic similarity, BLEU, and METEOR metrics - DrugChat substantially outperformed GPT-4.

**Fig. 3.**
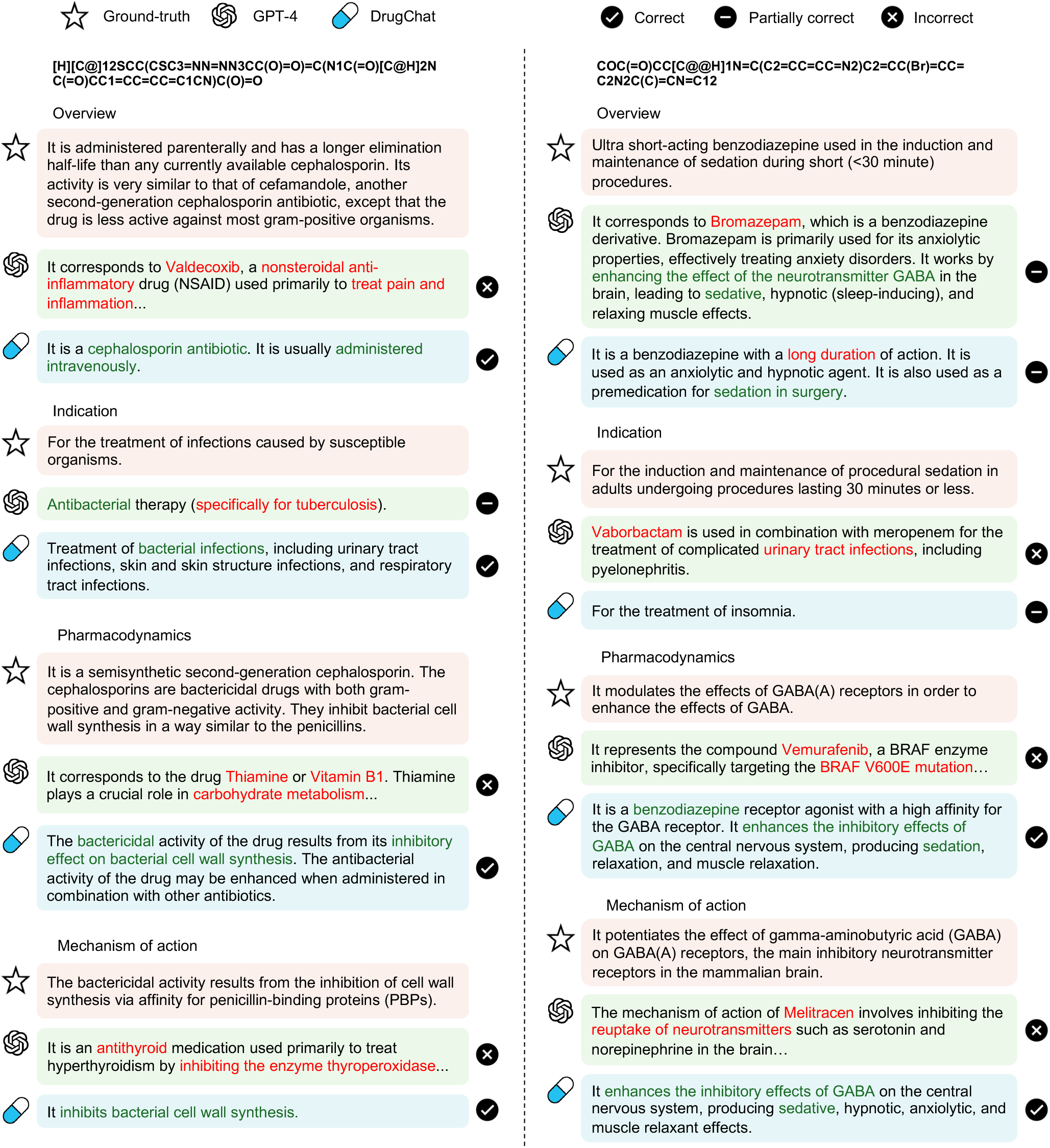
Examples of predictions generated by DrugChat and GPT-4 demonstrate that DrugChat’s predictions are more accurate than GPT-4’s. Text highlighted in red indicates incorrect predictions, while text highlighted in green indicates correct predictions.

Fig. 3 presents a comparison of predictions made by DrugChat and GPT-4 for several drug molecules randomly selected from the test set, which were not seen by the models during training. The predictions generated by DrugChat were notably more accurate compared to those by GPT-4, as validated by human expert ratings. For example, when provided with the SMILES of the molecule shown on the left in Fig. 3, DrugChat accurately predicted the molecule’s overview, indication, pharmacodynamics, and mechanism of action. The predictions aligned closely with the ground truth. DrugChat correctly classified the molecule as a cephalosporin antibiotic, identified its administration route as intravenous, and recognized its role in treating bacterial infections across multiple organs. Furthermore, it accurately predicted the mechanism of action, noting that the molecule inhibits bacterial cell wall synthesis. In contrast, GPT-4 produced largely incorrect predictions, misidentifying the molecule as a nonsteroidal anti-inflammatory drug (NSAID) for pain and inflammation, and incorrectly attributing its mechanism of action to COX-2 inhibition. As another example, when given the SMILES representation of the molecule shown on the right in Fig. 3, DrugChat accurately identified the molecule as a benzodiazepine and recognized its use for sedation during surgery. Regarding pharmacodynamics, DrugChat correctly indicated that the molecule enhances the inhibitory effects of GABA in the central nervous system, leading to sedation and muscle relaxation. Interestingly, when asked about the indication, DrugChat suggested the molecule’s use for treating insomnia, likely because benzodiazepines are sometimes used for that condition. In contrast, GPT-4 misclassified the molecule as either bromazepam or vaborbactam, providing largely incorrect information on both the indication and pharmacodynamics. Additionally, DrugChat delivered consistent predictions across various queries about the same molecule, whereas GPT-4 produced conflicting predictions, especially regarding the drug’s indication and pharmacodynamics. A mistake by DrugChat was its prediction that the molecule has a long duration of action, while it actually has a short duration.

Besides human evaluation, we also employed automated evaluation metrics, including semantic similarity, BLEU scores (40), and METEOR scores (41). Semantic similarity (ranging from -1 to 1, with higher values indicating better alignment) assesses the semantic alignment between the predicted text and the ground truth by leveraging language model embeddings (Methods). BLEU and METEOR scores quantify the overlap of words between the predicted and ground truth texts (Methods). Both BLEU and METEOR scores range from 0 to 1, with higher values indicating better performance. DrugChat demonstrated a large performance improvement over GPT-4 (Fig. 2d), achieving an average score of 0.47 in semantic similarity (compared to GPT-4’s 0.32), 0.3 in BLEU (compared to GPT-4’s 0.17), and 0.23 in METEOR (compared to GPT-4’s 0.14). These averages were calculated across predictions for drug indication, pharmacodynamics, mechanism of action, and overview.

DrugChat’s superior performance over GPT-4 is rooted in its molecule-aware architecture, explicitly designed for the complex domain of compound molecules. While GPT-4 excels as a general-domain textual language model, it has major limitations when applied to molecular understanding. These limitations arise from its processing approach, which interprets molecules’ SMILES representations merely as sequences of characters, devoid of chemical and spatial context. This hinders its ability to grasp the intricate structural relationships within molecules, which are crucial for accurate predictions of drug indications, pharmacodynamics, and mechanisms of action. In contrast, DrugChat is meticulously designed with dedicated molecular encoders, including a graph neural network (GNN) and a convolutional neural network (CNN), pretrained on large-scale molecule datasets. The GNN excels at modeling the connectivity and interactions within molecules, capturing the relational information vital for understanding molecular behavior. Meanwhile, the CNN adeptly recognizes patterns within molecular structures, similar to how it identifies objects from images. By leveraging these specialized encoders, DrugChat effectively discerns subtle yet significant molecular features that often elude more general models like GPT-4. This dual-encoder approach, combining the strengths of GNN and CNN, enables DrugChat to deliver superior predictions.

### DrugChat accurately predicts drug properties represented as discrete categories

Beyond generating detailed free-form predictions, DrugChat is also capable of predicting drug properties represented as discrete categories. Specifically, we focused on predicting molecules’ cytotoxicity to human cells, administration routes (e.g., oral, parenteral, topical), and their potential to function as prodrugs. To assess DrugChat’s cytotoxicity prediction performance, we used a test set of 190 compounds from the dataset curated by Wong et al. (21). For evaluating its predictions on administration routes and prodrug status, we randomly selected 385 molecules from the ChEMBL test set. None of these molecules were included in the training data. We compared DrugChat against several state-of-the-art (SOTA) LLMs, including GPT-4 (27), LLaMa (17), ChatGLM (42), and FastChat-T5 (43), all of which, like DrugChat, used the SMILES of molecules as input. Additionally, we benchmarked its performance against ImageMol (24), a SOTA specialized model designed for categorical molecular property prediction (Methods). Performance was assessed using macro-averaged F1 score (higher is better), which accounts for the imbalance in category labels (Extended Data Fig. 3).

Accurately predicting the cytotoxicity of molecules to human cells presents a significant challenge, as it requires a deep understanding of molecular properties and their interactions within cellular environments. We performed cytotoxicity predictions on three types of human cells: human liver carcinoma cells (HepG2), primary skeletal muscle cells (HSkMC), and human lung fibroblast cells (IMR-90). HepG2 cells are commonly used for studying hepatotoxicity and general cytotoxicity, while HSkMCs and IMR-90 cells offer potential advantages over immortalized cell lines like HepG2 for studying in vivo toxicity (21). For predicting cytotoxicity to HepG2, we used a prompt: ‘Is the molecule cytotoxic to human liver carcinoma cells (HepG2)? Please answer Yes or No.’ Similar prompts were used for the other classification tasks. DrugChat achieved F1 scores ranging from 0.57 to 0.71, significantly outperforming LLM baselines, which ranged between 0.17 and 0.5 (Fig. 4a). The task-specific ImageMol model performed significantly worse than DrugChat, with F1 scores between 0.36 and 0.43.

**Fig. 4.**
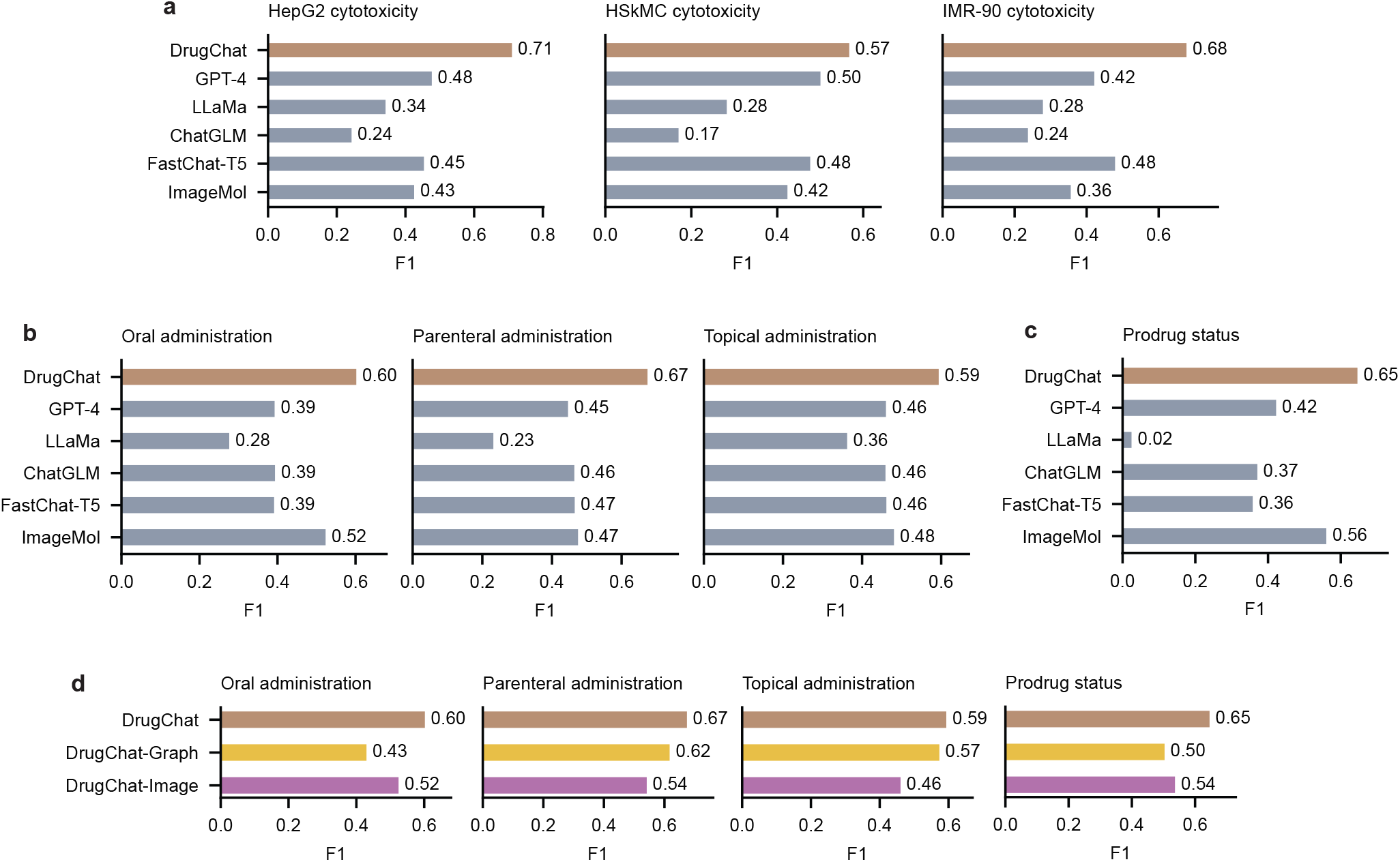
DrugChat accurately predicts drug properties represented as discrete categories. **a-c**, In tasks of predicting human cell cytotoxicity (**a**), administration routes (**b**), and prodrug status (**c**), DrugChat achieves significantly higher F1 scores compared to leading LLMs and specialized classifiers. **d**, DrugChat’s integration of molecular graph and image modalities outperformed the variants that rely on either graph or image modality alone.

Next, we focused on predicting the administration routes of molecules, a highly complex task that requires a deep understanding of the molecule’s physicochemical properties, its interactions with biological tissues, and its pharmacokinetic profile. Since a single drug molecule can be administered through multiple routes - for example, amoxicillin can be given orally or intravenously - we instructed DrugChat to predict a binary ‘yes/no’ for each route type, rather than limiting it to selecting only one option from all possible routes. DrugChat achieved F1 scores between 0.59 and 0.67, substantially surpassing baseline LLMs, which had F1 scores ranging from 0.23 to 0.47 (Fig. 4b). DrugChat also exceeded the performance of the task-specific ImageMol model significantly, which recorded F1 scores between 0.47 and 0.52.

Lastly, we predicted the potential of a molecule to act as a prodrug. A prodrug is a compound that, after administration, undergoes metabolic conversion within the body to become a pharmacologically active drug. Unlike active drugs, which exert their effects immediately upon administration, prodrugs are initially inactive or less active and require chemical transformation by the body’s metabolic processes to release their therapeutic potential. DrugChat achieved an F1 score of 0.65 (Fig. 4c), significantly out-performing baselines including GPT-4 (F1 score of 0.42), LLaMa (0.02), ChatGLM (0.37), FastChat-T5 (0.36), and ImageMol (0.56).

### DrugChat enables dynamic, iterative exploration of drug mechanisms and properties

DrugChat, as a multi-modal LLM, facilitates multi-turn interactions with users regarding the same molecule. The user initiates the conversation with a question, to which DrugChat generates a response. Based on this response, the user can ask follow-up questions, enabling an iterative dialogue process. Fig. 5 illustrates a conversation between DrugChat and a human user, where DrugChat provides accurate and detailed responses regarding the indication and pharmacodynamics of the input molecule. When the user inquired about the molecule’s indication, DrugChat accurately responded that it helps control blood sugar in individuals with type 2 diabetes by stimulating the pancreas to produce more insulin. Following this, the user asked what type of cells the molecule stimulates to release insulin, and DrugChat correctly identified pancreatic beta cells. The user then inquired about the molecule’s side effects and how to manage them, and DrugChat responded with appropriate information. Lastly, when asked about the site of the molecule’s metabolism, DrugChat accurately responded that it occurs in the liver. Through interactive dialogue with DrugChat, the user was able to achieve a more in-depth exploration of the molecule, uncovering increasingly detailed and nuanced information. DrugChat consistently understood the user’s intent and provided precise responses.

**Fig. 5.**
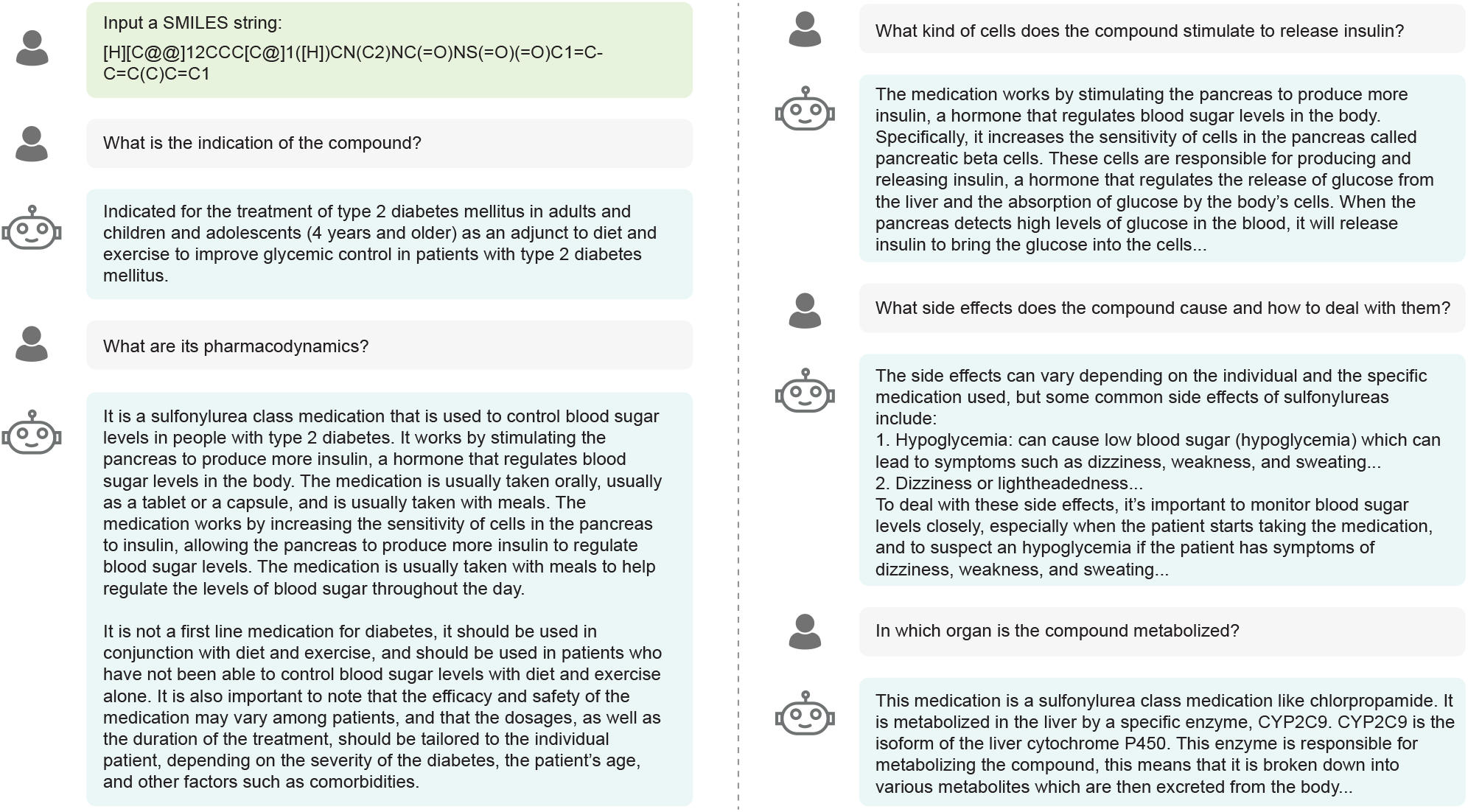
An exemplar multi-turn dialogue between DrugChat and a user regarding the same molecule.

### DrugChat’s integration of molecular graph and image modalities surpasses single-modality variants

To assess the impact of different molecule encoders on DrugChat’s performance, we developed two additional variants: DrugChat-Graph, which exclusively utilized molecular representations derived from the molecular graph via the graph neural network, and DrugChat-Image, which relied solely on representations extracted from the molecular image using the convolutional neural network (Methods). We compared the performance of these variants against the original DrugChat, which integrates representations extracted from both the image and graph. The original DrugChat consistently outperformed both variants across multiple prediction tasks (Fig. 4d), which demonstrates that the combined use of both image and graph representations is superior to relying on either modality alone. This can be attributed to the complementary nature of these modalities in capturing different aspects of molecule information. The molecular graph captures the topological relationships between atoms, which are crucial for understanding the molecule’s intrinsic properties, such as bond connectivity, electronic structure, and chemical reactivity. On the other hand, the molecular image highlights spatial patterns and visual features that are often critical in understanding the geometric and stereochemical properties of the molecule. These features are particularly relevant in identifying functional groups or understanding spatial interactions between atoms, which can influence molecular behavior and interactions with biological targets. By integrating both modalities, DrugChat benefits from the rich, multi-faceted information that each modality offers, leading to a more comprehensive and nuanced understanding of the molecule. In contrast, relying on a single modality may overlook critical information, leading to less effective predictions and a narrower understanding of the molecule’s characteristics.

## Discussion

DrugChat utilizes a single, unified framework to address a wide range of prediction tasks associated with drug discovery and development, including predicting indications, pharmacodynamics, mechanisms of action, cytotoxicity, administration routes, and so on. This is achieved simply by altering the prompts, thereby eliminating the need for model retraining. This approach significantly reduces the complexity associated with developing and maintaining a suite of specialized models, making it particularly beneficial in environments where resources for model development and maintenance are limited, or where rapid iteration and deployment are crucial. By enabling the model to extract and integrate knowledge across various domains, DrugChat can potentially identify patterns and relationships that might be missed when using isolated models. This cross-domain learning enhances the model’s ability to generalize across different tasks, leading to more robust and accurate predictions. This is evidenced by its superior performance compared to specialized models like ImageMol (Fig. 4a-c), which, due to their narrower focus and limited capacity to integrate cross-domain knowledge, may overlook certain patterns, leading to less optimal predictions.

As a multi-modal LLM, DrugChat generates detailed, human-like textual predictions across a wide array of complex drug-related aspects, including indications, mechanisms of action, and pharmacodynamics (Fig. 3). This marks an important shift from traditional models that often oversimplify the intricate and multifaceted nature of drug mechanisms and properties into predetermined categories.

For instance, a drug’s mechanism of action is rarely a single, straightforward pathway but rather involves a series of interconnected biological processes that can vary depending on the context. Categorical representations may obscure these layers of complexity, leading to a less nuanced understanding. In contrast, DrugChat’s free-form predictions naturally and comprehensively articulate these complexities, capturing subtle interactions, rare side effects, and context-dependent variations. This richness of detail ensures that DrugChat can support more informed decision-making in drug development, personalized medicine, and clinical practice. By generating predictions that mirror the narrative style found in scientific literature and clinical discussions, DrugChat aligns more closely with how healthcare professionals and researchers conceptualize and communicate drug information.

Unlike traditional models which typically provide static, one-off predictions, DrugChat can dynamically respond to a sequence of user queries, enabling users to uncover insights that might be missed in a single-pass analysis and allowing for a deeper investigation into various aspects of a molecule in question without needing to input new data or switch between different models. Each query builds on the previous ones, with DrugChat using the context established in earlier exchanges to refine and enhance its subsequent predictions. This interactive approach not only makes DrugChat more user-friendly but also aligns with the iterative nature of scientific inquiry. Moreover, this functionality allows for collaborative research efforts, where multiple stakeholders can engage with DrugChat over time, refining and expanding the analysis as new questions and data emerge.

One challenge with DrugChat lies in the interpretability of its predictions. While the model excels at generating natural language outputs that facilitate user interaction, the decision-making process behind these predictions remains largely opaque, typical of the black-box nature of LLMs. This opacity can become problematic in critical healthcare contexts where understanding the model’s reasoning is essential, particularly for tasks like drug mechanism predictions or risk assessments, which require a high degree of trust and transparency. Without clear insight into how the model reaches its conclusions, it can be difficult to validate or scrutinize its outputs. To address this, our future work will focus on developing more interpretable LLMs and integrating advanced explainability techniques.

The development of DrugChat opens up numerous promising research directions. One key area for future work involves enhancing the model’s accuracy and consistency by incorporating more diverse training datasets, especially those that represent underexplored drug categories. Additionally, integrating DrugChat with complementary computational tools, such as molecular docking simulations or quantum mechanical calculations, could yield a more holistic and synergistic approach to drug discovery. Exploring the application of DrugChat in real-world drug discovery pipelines is another exciting direction, with potential uses in predicting drug-drug interactions, optimizing formulations, and identifying off-target effects. Moreover, expanding the model’s capability to handle larger molecule structures, such as biologics, could significantly increase its utility and impact within the pharmaceutical industry.

## Methods

### Data collection and processing

We curated training data for DrugChat from public compound molecule databases, including ChEMBL, PubChem, and DrugBank. The ChEMBL database^1^ contains information for 2,354,965 chemical compounds. We downloaded the SQLite version of the dataset, last updated on February 28, 2023. From the full dataset, we collected 14,816 compounds that include drug-related information. The PubChem database^2^ contains information of 66,469,244 chemical compounds. We used the data version last updated on May 9, 2023. Of these compounds, 19,319 include drug-related information. The DrugBank database^3^ (version 5.1.10, released on January 4, 2023) contains 16,428 drug entries. From this, we selected 11,583 entries with available SMILES strings, all classified as small molecules, excluding those categorized as biotech. After further filtering to remove entries lacking annotations for drug indications, pharmacodynamics, or mechanisms of action, we retained 3,700, 6,649, and 5,846 drug molecules from ChEMBL, PubChem, and DrugBank, respectively. For each drug, we collected its SMILES string along with various attributes including free-form attributes such as indications, pharmacodynamics, and mechanisms of action, as well as categorical attributes like administration routes and prodrug status.

Using these drugs and their annotated attributes, we curated the training data for DrugChat. For each attribute *a* of a drug molecule *m*, we created a triplet consisting of the molecule’s SMILES representation, a textual prompt querying the value of *a*, and the corresponding ground truth value for *a*. Each attribute type had its own tailored prompt. For instance, for the attribute ‘drug indication’, the corresponding prompt is ‘what is its indication?’. The answer would be a textual description of the molecule’s indication, provided by human experts in the ChEMBL, PubChem, and DrugBank databases. As another example, for the attribute ‘prodrug status’, the corresponding prompt is: ‘can it potentially be used as a prodrug? Please answer Yes or No.’ The ground truth answer is ‘Yes’ if the molecule is a prodrug, and ‘No’ otherwise. In total, we curated 91,365 triplets.

### Model architecture

DrugChat is a multi-modal model that integrates information from three distinct modalities: graphs, images, and text. It consists of a graph neural network (GNN), a convolutional neural network (CNN), and a large language model (LLM). For a given molecule, its SMILES string is efficiently converted into a molecular graph and a molecular image using the RDKit software^4^.

In the molecular graph, nodes correspond to the molecule’s atoms, while edges represent the chemical bonds between them. Each atom is defined by its atom type and chirality, with 120 atom types in total, including a special ‘Unknown’ category for unidentified atoms. Atom chirality is categorized into four types: tetrahedral clockwise, tetrahedral counter-clockwise, unrecognized, and other forms. These atom type and chirality attributes serve as the initial features for each node. Chemical bonds are characterized by type and direction, with bond types classified as single, double, triple, or aromatic, and bond directions as none, end upright, or end downright. These bond attributes are used as initial features for each edge. All node and edge features are categorical, with each category encoded as a vector with learnable parameters. The molecular graph is input into a pretrained GNN, specifically a graph convolutional network (GCN) (15), to learn a representation vector for the entire graph. The GNN leverages the graph’s connectivity, along with the initial features of nodes and edges, to learn multiple layers of representations for each node. Using a neighborhood aggregation approach, the GNN iteratively updates a node’s representation vector by aggregating the representations of its neighboring nodes and edges (15). Thus, after *K* layers of representation learning in the GNN, information from nodes and edges is propagated through *K*-hop paths across the graph. To obtain the feature vector for the entire graph, an average pooling operation is applied to compute the mean of the representation vectors for all nodes after *K* layers of learning. The GNN in DrugChat consists of five layers, with approximately 0.5 million parameters. Both node and edge representation vectors have a dimensionality of 300. The GNN was pretrained using a self-supervised learning approach called context prediction (25), where the model learns to predict a molecule’s sub-graphs based on its neighboring sub-graphs (i.e., context). This pretraining was performed using 2 million unlabeled molecules from the ZINC15 database (28).

For the molecular image, we employed a pretrained convolutional neural network (CNN) to extract a representation vector. The CNN processes the input image through multiple layers of 2D convolution, where each layer applies convolutional filters to detect specific patterns in the image or in the feature maps from the previous layer. We used a pretrained ResNet-18 (16) model, specifically ImageMol (24), which consists of 18 convolution layers with a total of 11 million parameters. A global average pooling layer converts the output of the last convolution layer to a molecular image representation vector with a dimensionality of 512. ImageMol was pretrained on 10 million unlabeled images of drug-like, bioactive molecules from the PubChem database using self-supervised learning (SSL) techniques, such as molecular image reconstruction and contrastive learning. These SSL algorithms enabled ImageMol to map structurally similar molecules to nearby vectors in the embedding space, allowing the model to learn molecular properties at scale without requiring human-labeled data.

After extracting representation vectors from the molecule using the GNN and CNN, we apply two separate multilayer perceptrons (MLPs), referred to as the *adapters*, to transform these molecular representations into a format compatible with the LLM. LLMs typically use Transformer decoders (36) to model natural language as a sequence of tokens, where each token is represented as a vector (45). In DrugChat, the transformed molecular representations are treated as tokens and appended to the language token sequence, which was converted from the input prompt.

This combined sequence is then passed into the LLM, which uses multi-head self-attention mechanisms (36) to generate new tokens. These generated tokens form the final prediction. DrugChat uses Vicuna-13B (35) as the LLM, which contains 13 billion parameters. It was fine-tuned from Llama-13B (17) on a dataset of 70K user-shared dialogues collected from ShareGPT.com (containing conversations between human users and ChatGPT). It retains the same architecture as Llama-13B, including 40 Transformer layers with 40 attention heads and an embedding dimension of 5120. Llama-13B was pretrained on a multi-terabyte text corpus, including Wikipedia and various other sources gathered from the Internet, to predict the next word based on the preceding context. The adapter responsible for converting molecular graph representations into LLM tokens consists of two layers, with an input dimension of 300, a hidden dimension of 5120, and an output dimension of 5120, totaling 28M parameters. Similarly, the adapter for converting molecular image representations includes two layers, with an input dimension of 512, a hidden dimension of 5120, and an output dimension of 5120, amounting to 29M parameters. Both MLPs use the GELU activation function (46) in the hidden layer.

For a target answer *T* that has *L* text tokens, DrugChat computes the probability of generating *T* as follows:

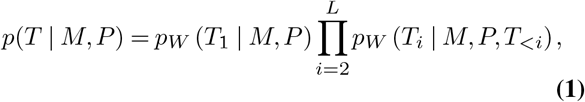

where *M* represents the input molecule and *P* is the input prompt. We denote the *i*-th token as *T*_*i*_ and all preceding tokens as *T*_*<i*_. Model parameters are denoted by *W*. The generated token sequence is compared to the ground truth tokens to compute the negative log likelihood (NLL). The parameters *W* are optimized by minimizing the sum of NLL over all training data.

### Model training

We trained the DrugChat model using the AdamW optimizer (47) with 10 epochs, employing default parameters *β*_1_ = 0.9, *β*_2_ = 0.999 and a weight decay of 0.01. In the first epoch, we applied a linear warmup strategy, gradually increasing the learning rate from 10^−6^ to 2 × 10^−5^, which was then reduced to 10^−6^ using a cosine decay scheduler over the remaining 9 epochs. The batch size was set to 16, and training was performed on NVIDIA A100 GPU with 80GB memory.

For training the specialized classification model ImageMol (24), we appended a classification head to the pretrained ResNet encoder (16). The classification head was a linear layer, producing an output vector with dimensions matching the number of classes in the prediction task. Given a molecular image, the pretrained encoder extracts a representation, which is subsequently passed to the classification head to predict the class label. The training data includes approximately 51,000 examples curated from 2,940 ChEMBL compounds, along with about 118,000 examples from 39,000 compounds in the dataset curated by Wong et al. (21). We optimized the model using a cross-entropy loss function and the AdamW optimizer for 50 epochs with a batch size of 128. The maximum learning rate was set to 10^−3^, while all other optimization settings followed those used for DrugChat.

### Prompts

The prompts used by DrugChat to predict cytotoxicity to HepG2, HSkMC, and IMR-90 were ‘Is the molecule cytotoxic to human liver carcinoma cells (HepG2)?’, ‘Is the molecule cytotoxic to primary skeletal muscle cells (HSkMC)?’, and ‘Is the molecule cytotoxic to human lung fibroblast cells (IMR-90)?’, followed by an instruction ‘Please answer Yes or No’. The prompts used by DrugChat to predict administration routes - oral, parenteral, and topical - were: ‘Is the molecule taken orally?’, ‘Is the molecule administered parenterally?’, and ‘Is the molecule applied topically?’, followed by an instruction ‘Please answer Yes or No’. For predicting drug indications, pharmacodynamics, and mechanisms of action, GPT-4 was prompted with: ‘Given the SMILES of a molecule: [a SMILES string]’, followed by questions like ‘What is its indication?’, ‘What are its pharmacodynamics?’, and ‘What is its mechanism of action?’. For predicting cytotoxicity, administration routes, and prodrug status, baseline LLMs used similar prompts: ‘Given the SMILES of a molecule: [a SMILES string]’, along with questions such as ‘Is the molecule cytotoxic to human liver carcinoma cells (HepG2)?’, ‘Is the molecule cytotoxic to primary skeletal muscle cells (HSkMC)?’, and ‘Is the molecule cytotoxic to human lung fibroblast cells (IMR-90)?’, ‘Is the molecule applied topically?’, ‘Is the molecule taken orally?’, ‘Is the molecule administered parenterally?’, and ‘Can it potentially be used as a prodrug?’, followed by the instruction ‘Please answer Yes or No’.

### Model evaluation

We evaluated the free-form predictions of drug indications, pharmacodynamics, and mechanisms of action using both human assessment and automated metrics.

#### Human evaluation

In the human evaluation, experts specializing in drug molecules assessed the model’s predictions using a 3-point Likert scale with an additional ‘Unknown’ option. The scales were defined as follows: 1) [Correct] — The prediction is mostly consistent with the ground truth or a subset of the ground truth, possibly extending it with additional plausible details; 2) [Partially Correct] — The prediction includes some correct descriptions but also introduces conflicting elements when compared to the ground truth or domain knowledge; and 3) [Incorrect] — The prediction is incorrect, irrelevant, or incomplete. Evaluators were asked to choose one of the three options and they did not know which model generated the predictions.

#### Automatic evaluation metrics

We conducted automatic evaluations using three metrics: semantic similarity,

BLEU (40), and METEOR (41) scores. Semantic similarity was calculated as the cosine similarity between the sentence embeddings of the ground-truth and model-predicted texts, with embeddings generated using a pretrained sentence Transformer model All-MiniLM-L6-v2 (48). Let SE represent the sentence embedding model. The embedding of the ground-truth text *t*_*g*_ is denoted as *e*_*g*_ = SE(*t*_*g*_), while the embedding of the predicted text *t*_*p*_ is *e*_*p*_ = SE(*t*_*p*_). The cosine similarity between these embeddings is defined as

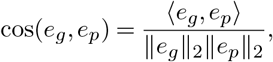

where ⟨*e*_*g*_, *e*_*p*_⟩ represents the dot product between the two vectors, and ∥*e*_*g*_∥_2_ denotes the L2 norm of the vector *e*_*g*_. With this sentence Transformer, the embeddings of semantically similar sentences are positioned closer in the embedding space, resulting in higher cosine similarity values. For BLEU scores (40), we used the BLEU-1 score without applying the brevity penalty (40). The BLEU-1 score is a special case of the general BLEU-n metric, which measures the modified precision of n-grams (sequences of *n* consecutive words). Let *ŷ* represent the predicted sentence, and *y* the ground-truth sentence. Define *G*_*n*_(*ŷ*) as the set of all n-grams in the predicted sentence. Let *C*(*s, y*) be an indicator function, which equals 1 if the n-gram *s* appears in *y*; otherwise, *C*(*s, y*) = 0. The BLEU-n score is then calculated as follows:

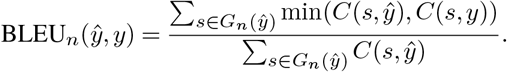

The METEOR score (41) is viewed as an enhancement over the BLEU score. It is calculated using the harmonic mean of unigram precision and recall, with greater emphasis on recall. Additionally, it accounts for stemming and synonym matching, in addition to exact word matching.

### DrugChat accurately predicts drug-related keywords

We extracted keywords related to drugs, diseases, and medical conditions from the ground-truth answers in the DrugBank test set using the Python library scispaCy (44). After extraction, we manually refined the keyword set by removing less relevant keywords, duplicates, and those appearing in fewer than three ground-truth answers. For each keyword in a ground-truth answer, scispaCy was used to identify semantically similar terms in the corresponding DrugChat-predicted text. If the similarity between the keyword and a predicted term exceeded 0.8 (on a scale of 0 to 1), we considered the term a match to the keyword. The matched terms were manually verified to ensure they were semantically similar to the corresponding keywords. We define recall for a keyword as the ratio of the number of matches to the keyword’s frequency. Extended Data Fig. 1 presents the recall and frequency of keywords.

DrugChat achieved a recall between 30% and 80% for the majority of keywords, despite the fact that most keywords occur fewer than ten times. This low frequency poses challenges for accurate prediction. However, DrugChat demonstrated strong recall in many cases, such as 75% for the keyword ‘respiratory depression’ and 69% for ‘hypertension’. Although DrugChat shows lower recall for some medical keywords, this does not necessarily indicate poor performance in predicting molecule functions related to those keywords. For example, while DrugChat incorrectly predicted that remimazolam is used to treat insomnia, the drug is actually used for the maintenance of sedation during short procedures, a function related to insomnia.

## Data availability

The dataset curated and used in this work is available at https://drive.google.com/drive/folders/1ofHOV5UFJUf2Xb-UljbK--wILF-_vBod?usp=sharing

## Code availability

The source code of this work is available at https://github.com/youweiliang/drugchat

## Acknowledgements

This work was supported by National Science Foundation IIS2405974 and National Institutes of Health 1R21GM154171-01, and was supported by the National Institutes of Health Bridge2AI program [1OT2OD032742 to T.I.].

---

**Extended Data Fig 1.**
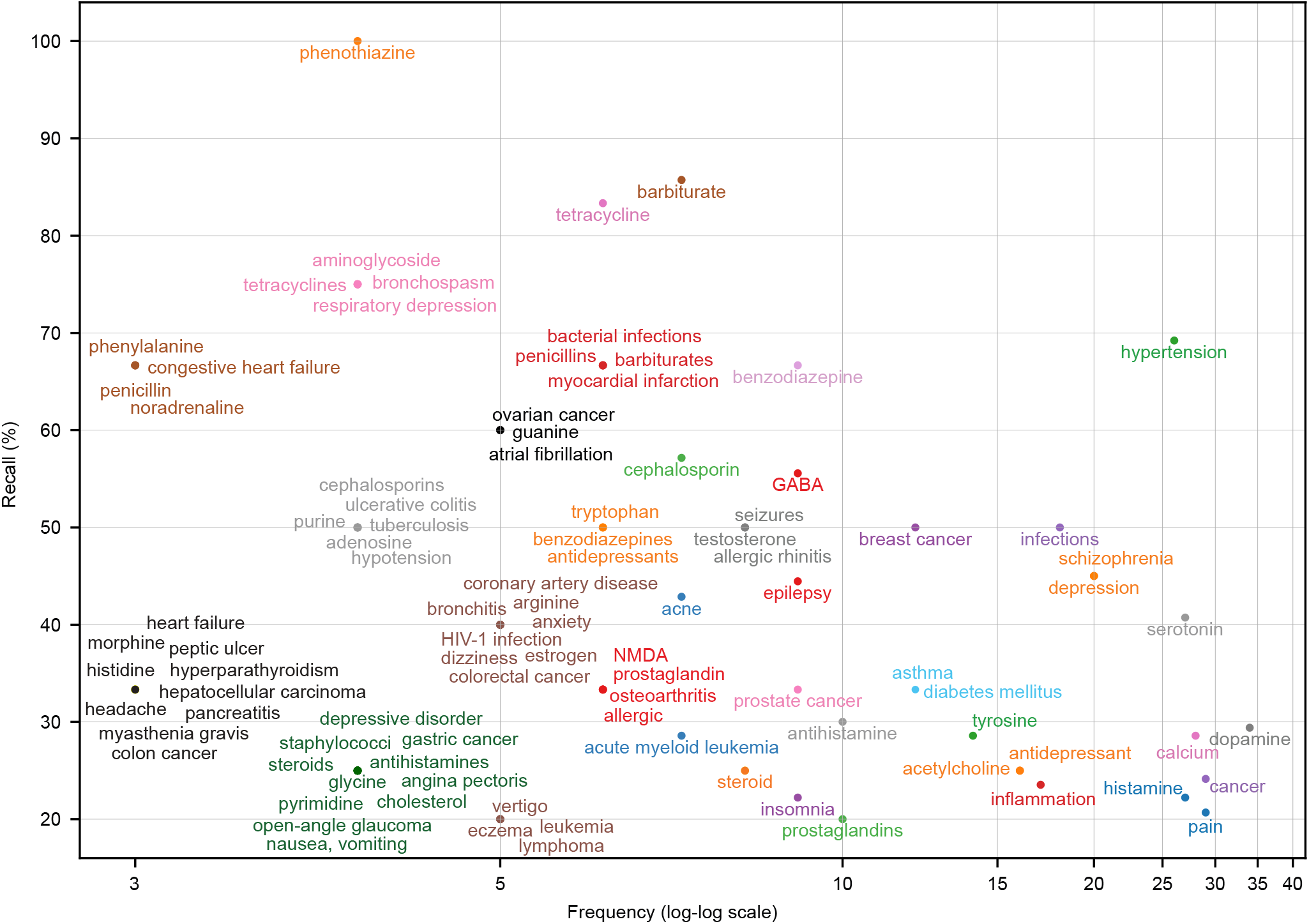
DrugChat achieved good recall rates in predicting keywords related to drugs, diseases, and medical conditions. The x-axis represents the frequency of these keywords.

**Extended Data Fig 2.**
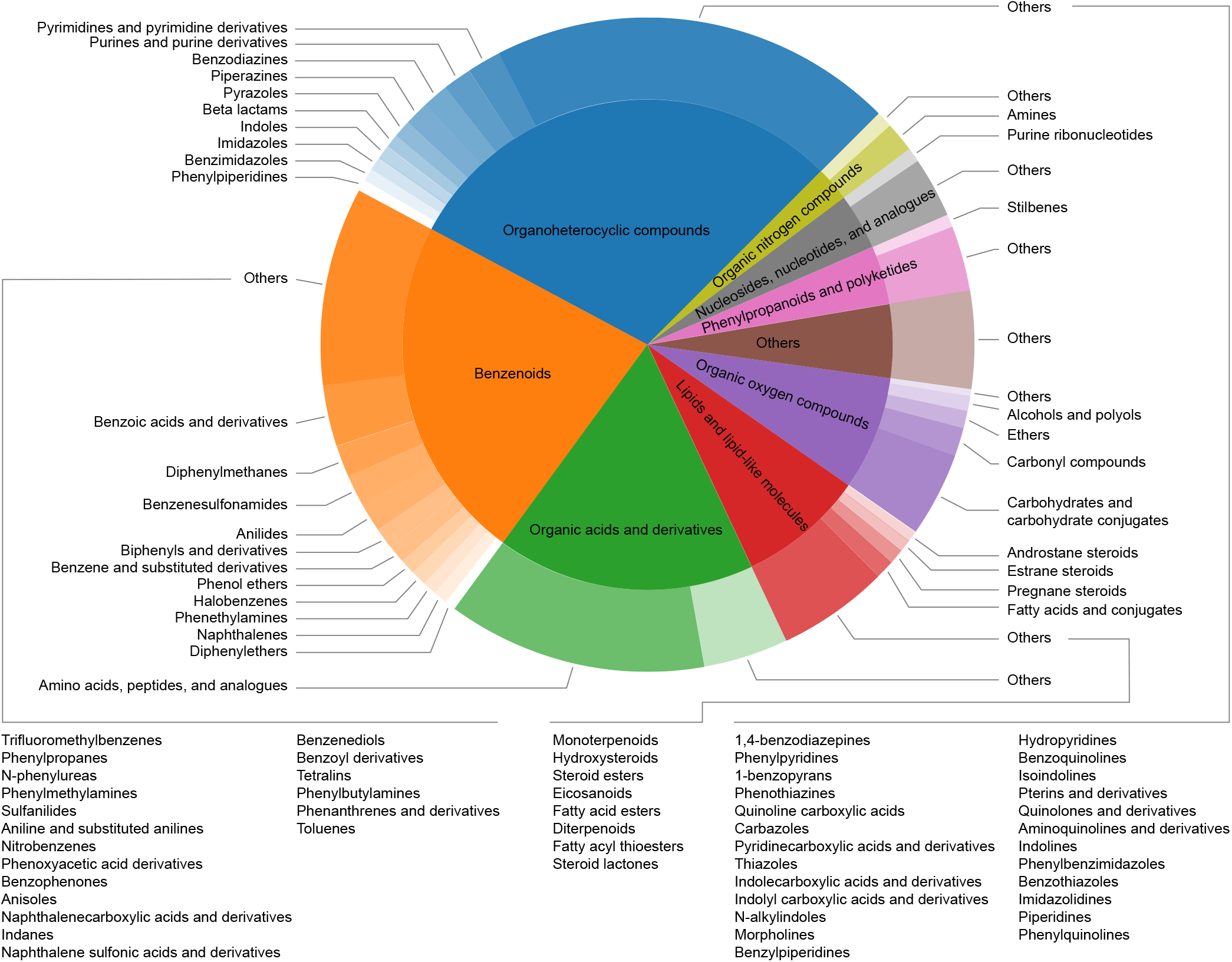
Our curated dataset features a diverse distribution of compound categories. The inner disk represents the compound superclasses, while the outer ring shows the corresponding subclasses.

**Extended Data Fig 3.**
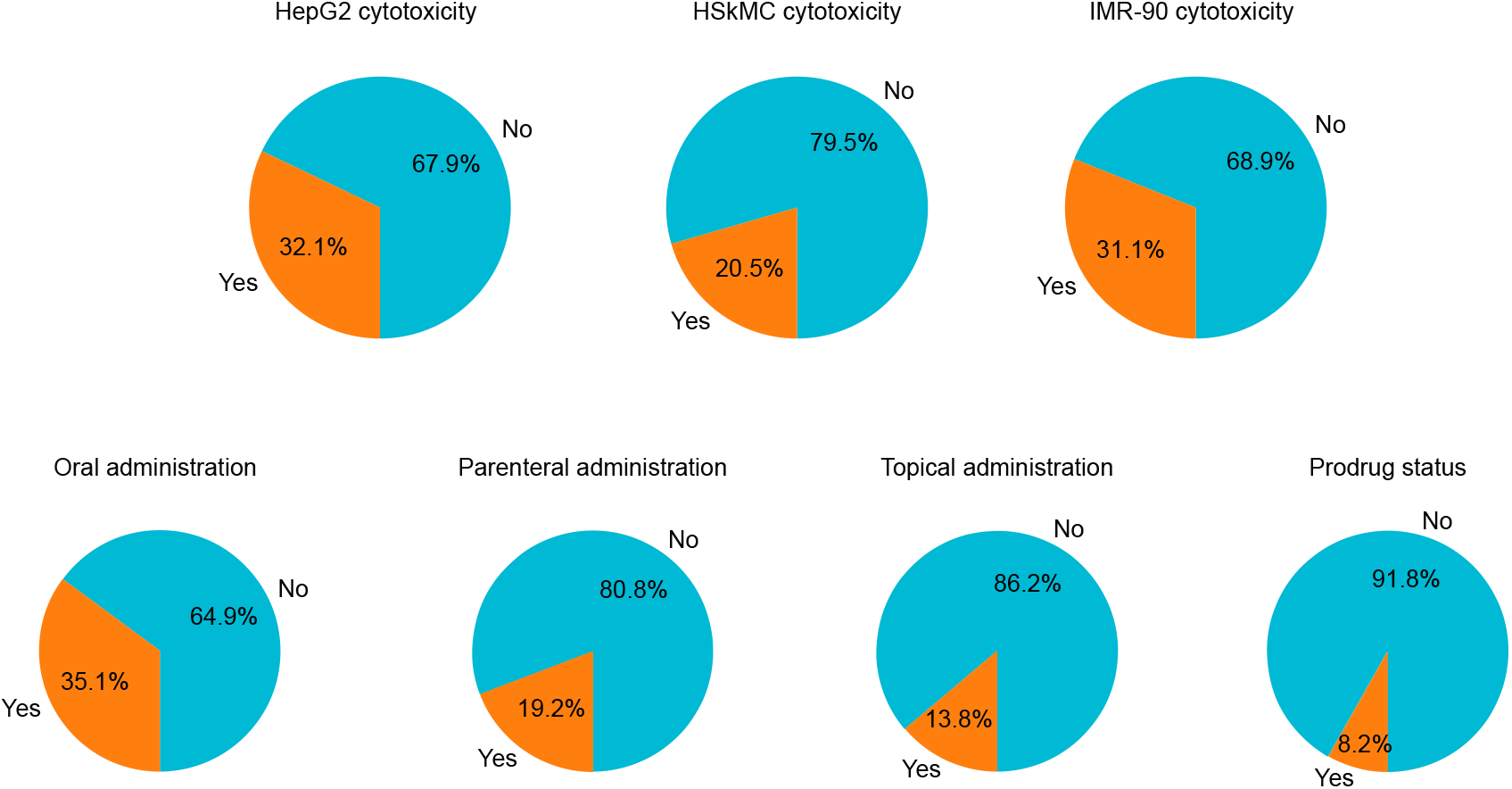
The distribution of ground-truth answers in the test set across the tasks of predicting cytotoxicity, administration routes, and prodrug status.

## Footnotes

1 https://www.ebi.ac.uk/chembl/

2 https://pubchem.ncbi.nlm.nih.gov/

3 https://go.drugbank.com/releases/latest

4 https://www.rdkit.org

